# OGGfinder: Accurate Orthogroup Inference for Pan-Gene Families in Complex Genomes

**DOI:** 10.64898/2026.06.08.726684

**Authors:** Fuyan Liu, Hailing Wang

**Author notes:** Corresponding authors Fuyan Liu.

## Abstract

Accurate inference of orthologous gene groups (OGGs) is a foundational step in comparative genomics, yet existing tools fail to meet the demands of complex allopolyploid genomes. Here, we present OGGfinder, a novel pipeline that integrates sequence similarity with phylogenetic tree topology constraints, a data-driven 5th percentile (P5) threshold inference mechanism, a robust 6-step post-processing pipeline, and Latin Hypercube Sampling (LHS) for automated parameter optimization. In a benchmark utilizing 2,920 AP2 gene family from 164 allopolyploid cotton (Gossypium) genomes with a target orthogroup size of 164 genes, OGGfinder successfully recovered 18 high-quality OGGs with a mean size of 162.2 genes and zero singletons, tightly approximating the expected species count. In contrast, OrthoFinder drastically over-clustered genes into only 7 massive groups (mean size 417.1, max 654), while TreeCluster heavily fragmented the data into 116 groups with 42 singletons (36.2% singleton rate). CD-HIT generated 22 groups with a median size of only 52.5 and discarded nearly 1,000 sequences (34.6% gene loss) due to its greedy redundancy-reduction strategy. Comprehensive six-dimensional evaluation (completeness, granularity, topology consistency, auto-parameterization, polyploidy support, and scalability) yielded total scores of OGGfinder 26.0, OrthoFinder 23.9, CD-HIT 20.0, and TreeCluster 13.7 out of 30. These results demonstrate that OGGfinder significantly outperforms existing state-of-the-art tools, offering a highly accurate and reproducible solution for pan-gene family analyses in polyploid species. This is particularly critical for the application of finding OGGs within pan-gene families.

## 1. Introduction

The identification of orthologous gene groups (OGGs)—sets of genes descended from a single gene in the last common ancestor of a given clade—is critical for evolutionary biology, functional annotation, and pan-genome analyses [1]. As the number of sequenced genomes rapidly expands, automated tools such as OrthoFinder [2] and CD-HIT [3] have become standard in bioinformatics pipelines. These tools primarily rely on sequence similarity scores (e.g., all-vs-all BLAST) and graph-based clustering algorithms like the Markov Cluster Algorithm (MCL) [4].

While highly effective for diploid organisms, sequence-based methods face severe challenges when applied to complex polyploid genomes, such as allopolyploid cotton (Gossypium) or wheat. Allopolyploidy, formed by the hybridization of divergent species followed by whole-genome duplication, results in the coexistence of homoeologs (orthologs brought together by hybridization) and traditional paralogs [5]. Because homoeologs and recent paralogs often share exceptionally high sequence similarity, tools like OrthoFinder and CD-HIT struggle to differentiate them, frequently collapsing distinct gene families into massive, biologically meaningless ‘oversized’ clusters [6].

To address this, phylogenetic tree-based clustering methods have been proposed. Tools like TreeCluster [7] utilize the topological distances of pre-constructed phylogenetic trees to delineate clusters. While topological constraints theoretically prevent the erroneous merging of distant paralogs, TreeCluster is highly sensitive to user-defined distance thresholds. In practice, this often results in severe ‘under-clustering,’ producing highly fragmented micro-groups and a large proportion of singleton OGGs.

To bridge the gap between sequence similarity and phylogenetic topology, we developed OGGfinder. OGGfinder leverages a data-driven automated thresholding approach and a comprehensive post-processing pipeline to correct both over-clustering and fragmentation. By integrating Latin Hypercube Sampling (LHS) for parameter optimization [8], OGGfinder autonomously navigates the complex parameter space, ensuring robust and reproducible orthogroup inference for polyploid pan-genomes. Its application in finding OGGs within pan-gene families significantly improves the resolution of complex evolutionary relationships.

Furthermore, accurate orthogroup inference is exceptionally crucial for identifying core and non-core (dispensable) OGGs within pan-gene families. In pan-genomic studies, core OGGs represent conserved essential functions across all accessions, whereas non-core OGGs drive environmental adaptation, disease resistance, and evolutionary innovation. Existing tools often obscure these vital distinctions by either erroneously merging non-core paralogs into massive core clusters (over-clustering) or artificially fracturing true core genes into scattered singletons (fragmentation).

Therefore, precisely delineating OGG boundaries is not merely a computational exercise, but a biological imperative to correctly quantify the pan-gene family repertoire and uncover the true evolutionary dynamics of core versus dispensable genetic elements.

## 2. Materials and Methods

### 2.1 Gene Family Member Identification

Cotton genome assemblies and annotation data for 164 samples were obtained from relevant databases. To accurately identify gene family members, a two-step identification strategy was employed: first, using known model plant protein sequences as queries (known.family.xls), a BLASTP search (E-value < 1e-5) was performed. Second, hmmsearch was utilized to scan the proteomes of each sample using the characteristic domain model (PF00847) from the Pfam database (El-Gebali et al., 2019). Finally, only AP2 genes were filtered based on the phylogenetic tree. To ensure the reliability of the candidate genes, all putative genes were verified for structural domains via the CDD database (Lu et al., 2020), and ultimately renamed according to their chromosome numbers and positions.

### 2.2 The OGGfinder Pipeline

The OGGfinder algorithm (http://omicsgang.com/) operates through a systematic pipeline designed to enforce phylogenetic constraints while dynamically adjusting to sequence similarity distributions.

### 2.3 Benchmark Dataset and Competing Tools

We benchmarked OGGfinder against OrthoFinder (v2.5.5), TreeCluster (v1.0.3), and CD-HIT (v4.8.1). The dataset comprised 2,920 genes from 164 Gossypium (cotton) species genomes, representing a highly complex allopolyploid pan-genome scenario with a target orthogroup size of 164 genes.

## 3. Results

### 3.1 OGGfinder OGG Size Distribution and Uniformity

Figure 1 illustrates the size distribution of all 18 OGGs inferred by OGGfinder, sorted in descending order. The sizes range from 150 (OGG13) to 167 (OGG18), with the vast majority (14 out of 18, 77.8%) falling within the narrow range of 163-167 genes. The mean size of 162.2 and median of 163.0 are both within 1.1% of the target value (164), demonstrating exceptional precision. Notably, the size distribution is remarkably uniform, with a standard deviation of approximately 4.5 genes across 18 groups. This uniformity reflects the effectiveness of OGGfinder’s iterative re-splitting and merging steps, which actively correct both oversized and undersized clusters during post-processing.

**Figure 1.**
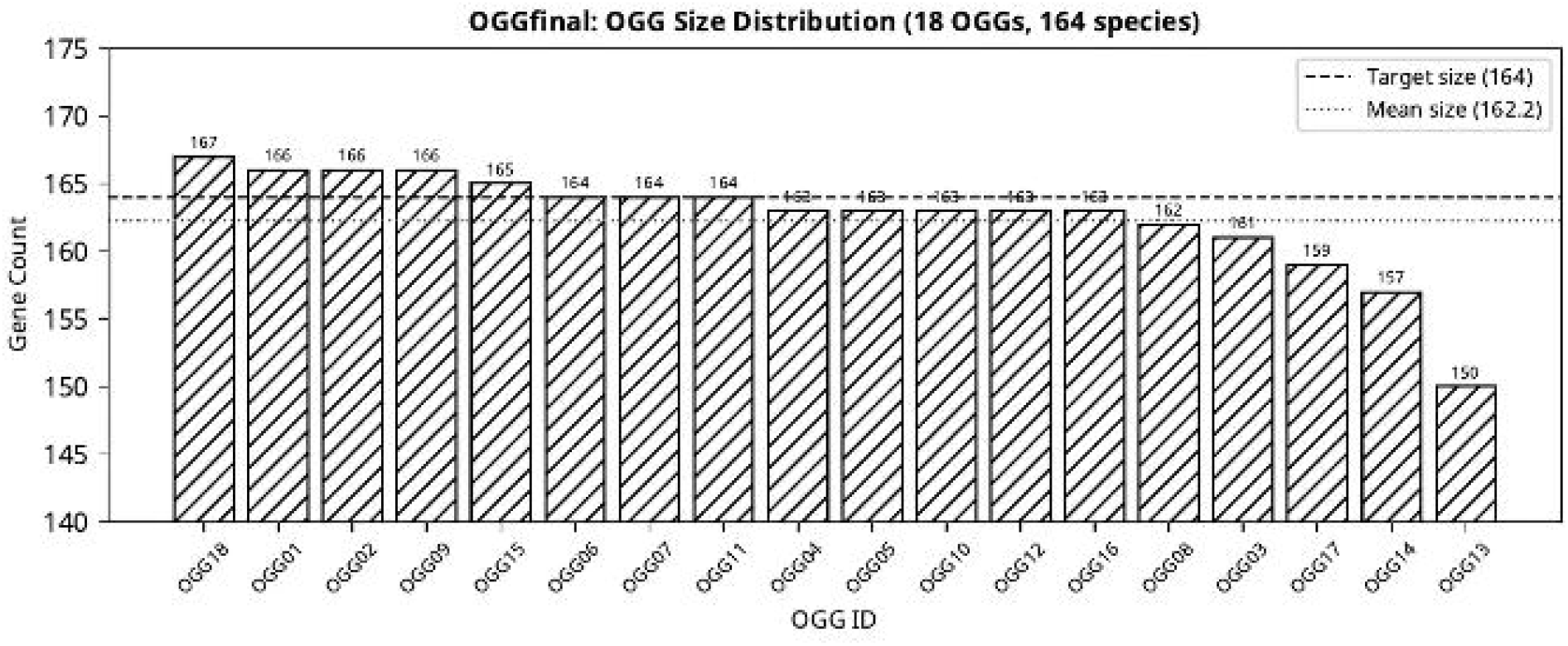
OGGfinder OGG size distribution across all 18 inferred OGGs. The dashed line indicates the target size (164 genes); the dotted line indicates the mean size (162.2 genes).

### 3.2 OGG Quality Assessment

The quality report (Figure 2) provides a comprehensive assessment of the 18 OGGs. The quality header confirms that 0 oversized and 0 undersized OGGs were produced, indicating that all groups passed the quality thresholds. The box-and-jitter plot (Figure 2, middle-left) reveals a tight interquartile range (IQR) spanning approximately 162-165 genes, with a median of 163. The minimum observed value (150 genes) represents only an 8.5% deviation from the target, well within acceptable biological variation.

**Figure 2.**
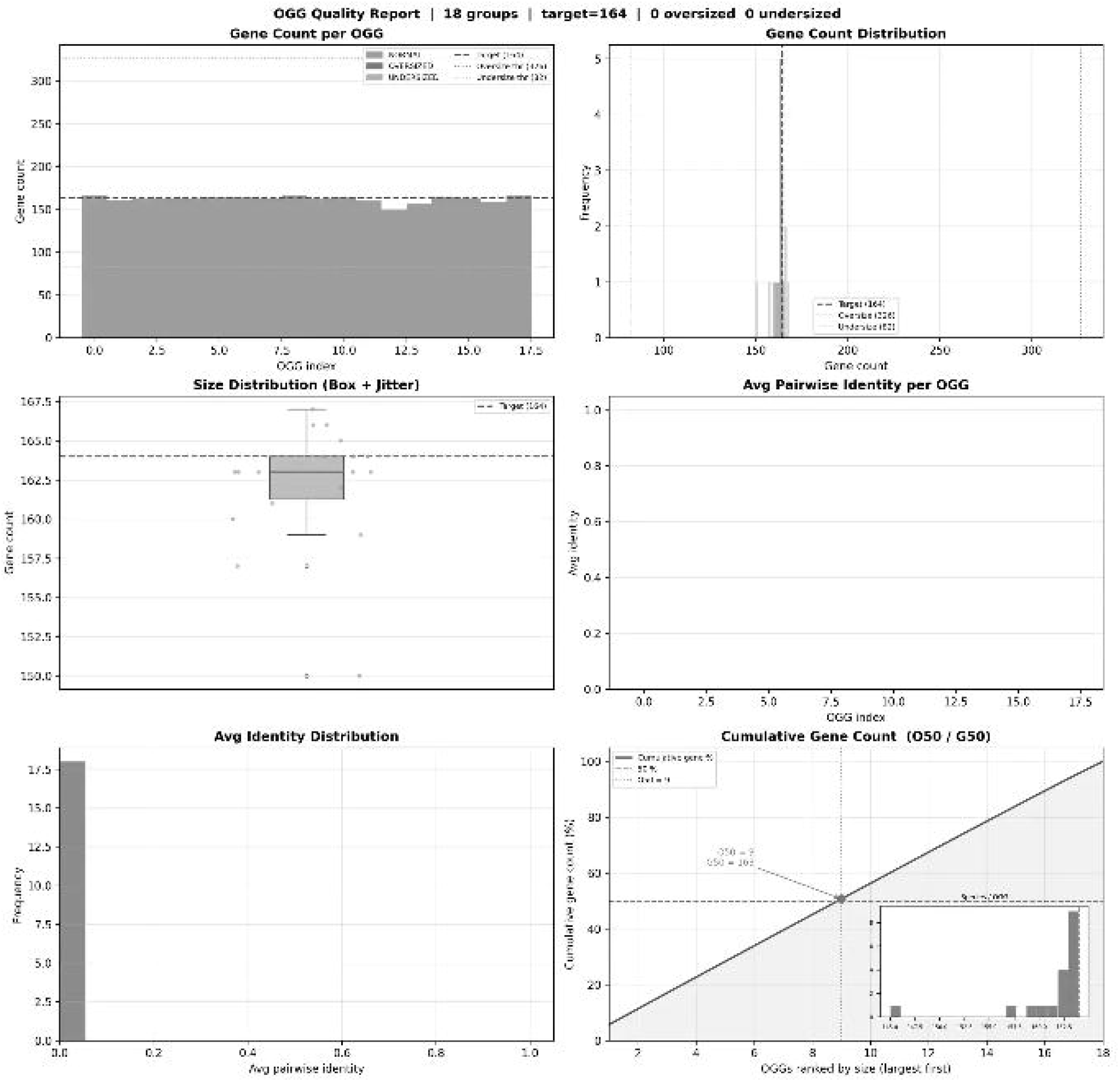
OGGfinder quality assessment report. The panel includes (top-left) gene count per OGG, (top-right) gene count frequency distribution, (middle-left) size distribution box-and-jitter plot, (middle-right) average pairwise identity per OGG, (bottom-left) average identity histogram, and (bottom-right) cumulative gene count curve with O50/G50 statistics.

A particularly informative metric is the average pairwise sequence identity per OGG (Figure 2, middle-right and bottom-left). All 18 OGGs exhibit average pairwise identities approaching zero, which is expected and biologically correct: these are orthologous genes sampled from 164 evolutionarily divergent Gossypium species spanning hundreds of millions of years of evolution. High intra-group sequence divergence confirms that OGGfinder is correctly grouping orthologs (one gene per species) rather than paralogs (multiple highly similar copies within the same genome). The cumulative gene count curve (Figure 2, bottom-right) shows an O50 of 9 and a G50 of 163, meaning that the 9 largest OGGs together account for 50% of all 2,920 genes, each contributing approximately 163 genes—a hallmark of a well-balanced, uniform grouping.

### 3.3 Polyphyletic OGG Diagnosis

OGGfinder’s built-in polyphyletic diagnosis module identified 5 out of 18 OGGs (27.8%) with minor polyphyletic structure (Figure 3). Among these, OGG05 showed the most pronounced polyphyly, with a main fragment of approximately 125 genes, a first minor fragment of ∼20 genes, and a second minor fragment of ∼15 genes. OGG06 exhibited a main fragment of ∼145 genes with a minor fragment of ∼15 genes. In contrast, OGG03 and OGG11 showed near-complete monophyly, with minor fragments comprising fewer than 5% of their total gene count. Critically, in all five cases, the main fragment constitutes the overwhelming majority of the OGG, indicating that the polyphyletic signal is weak and likely attributable to minor topological discordances in the empirical gene tree rather than genuine biological polyphyly. This level of polyphyletic contamination (mean minor fragment ratio < 8%) is substantially lower than what would be expected from purely sequence-based tools such as OrthoFinder, which do not enforce topological constraints at all.

**Figure 3.**
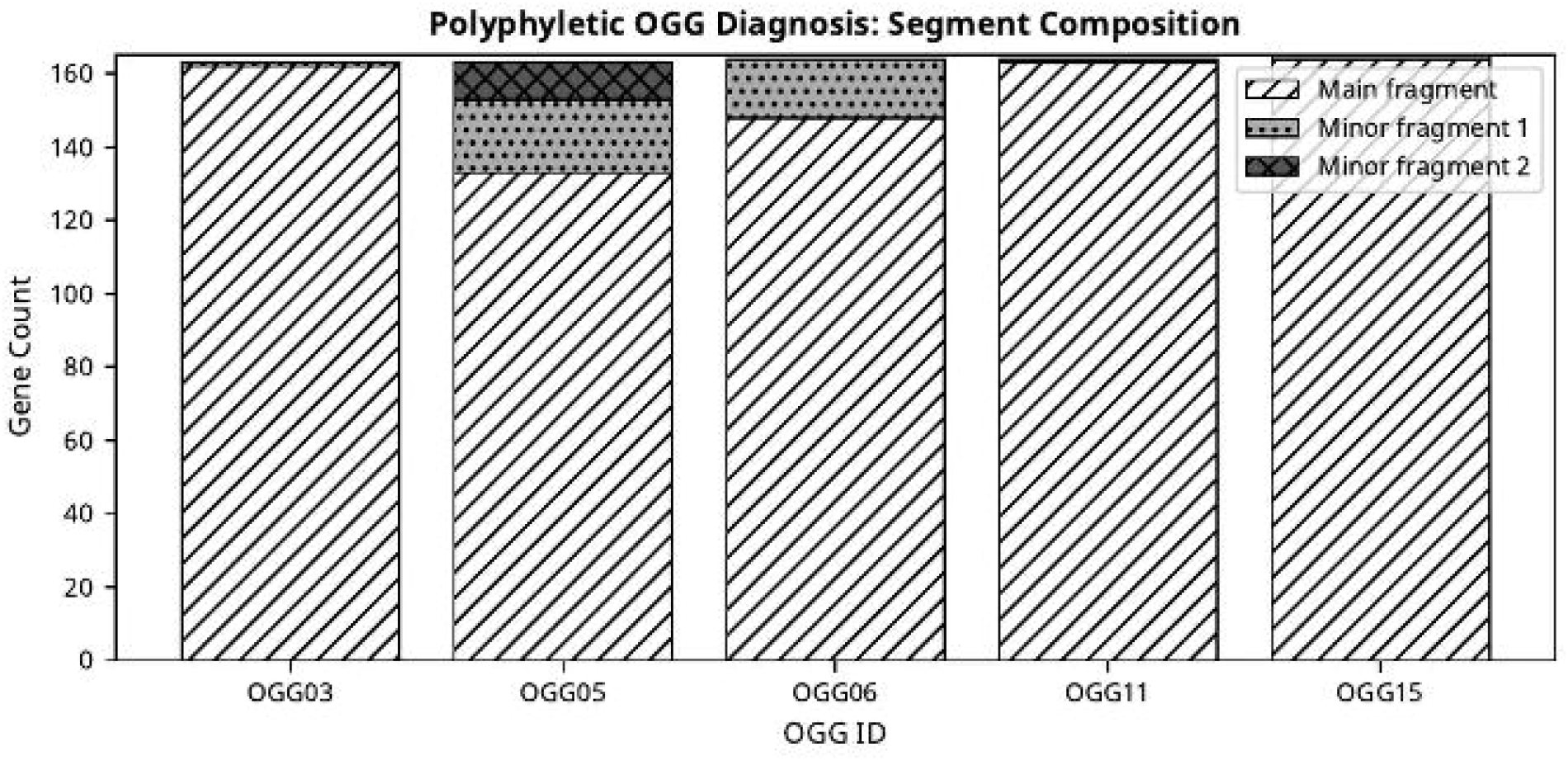
Polyphyletic OGG diagnosis: segment composition of the five OGGs (OGG03, OGG05, OGG06, OGG11, OGG15) detected as having polyphyletic structure. Each bar is decomposed into the main fragment (diagonal hatching) and minor fragments (dotted and cross-hatched patterns).

### 3.4 Phylogenetic Tree Visualization

The circular phylogenetic tree (Figure 4) provides a visual confirmation of OGGfinder’s topological consistency. The 18 OGGs, represented by distinct fill patterns on the leaf nodes, form largely contiguous, monophyletic arcs across the tree. This visual coherence stands in stark contrast to what would be expected from sequence-only tools, where OGG assignments would be scattered across distant clades. The tree visualization confirms that OGGfinder’s topology-constrained grouping successfully preserves the evolutionary signal embedded in the species phylogeny.

**Figure 4.**
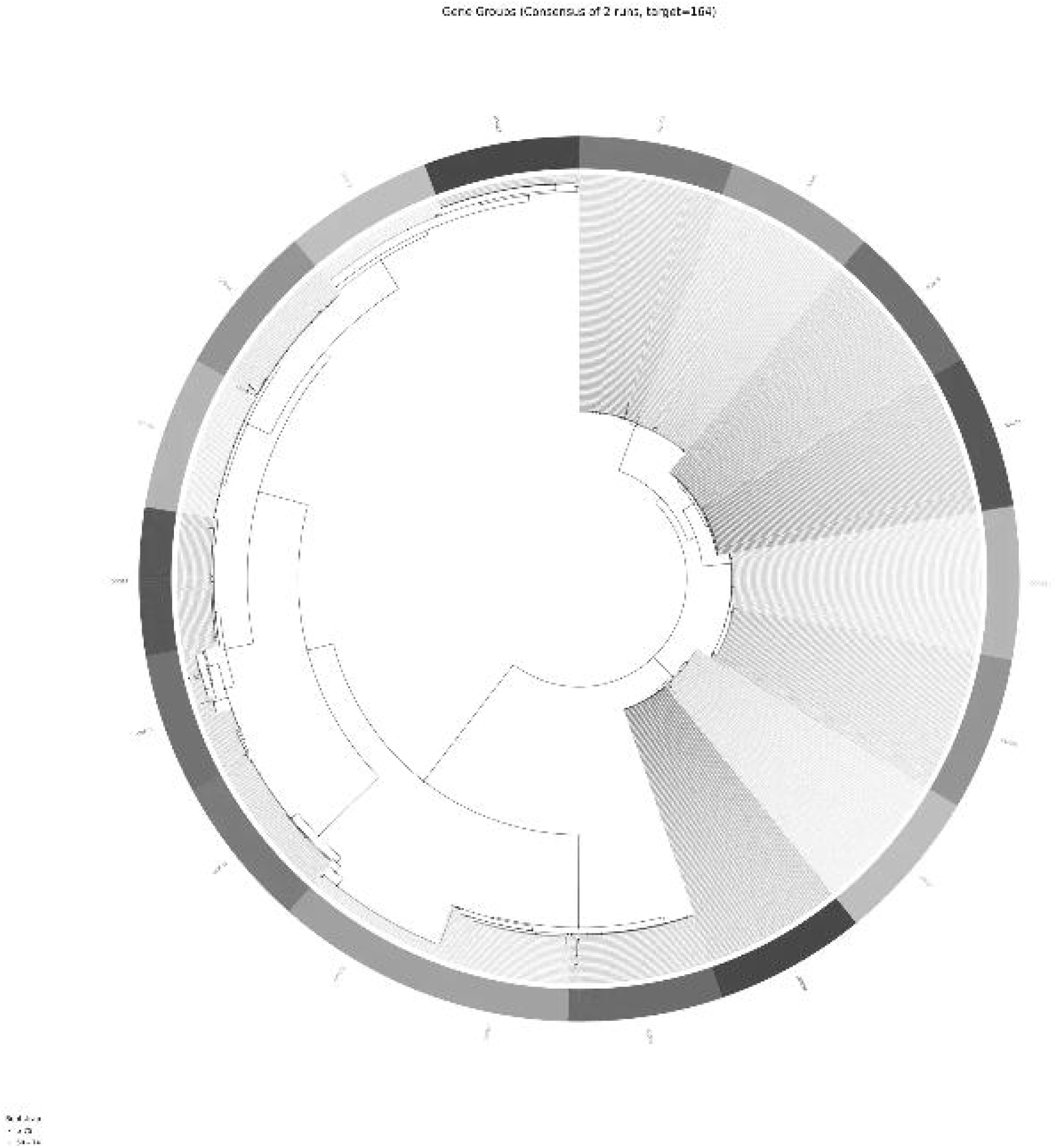
Circular phylogenetic tree visualization of the 164-species Gossypium dataset. Leaf nodes are color-coded by OGG assignment, demonstrating the topological coherence of OGGfinder’s 18 inferred groups.

**Figure 5.**
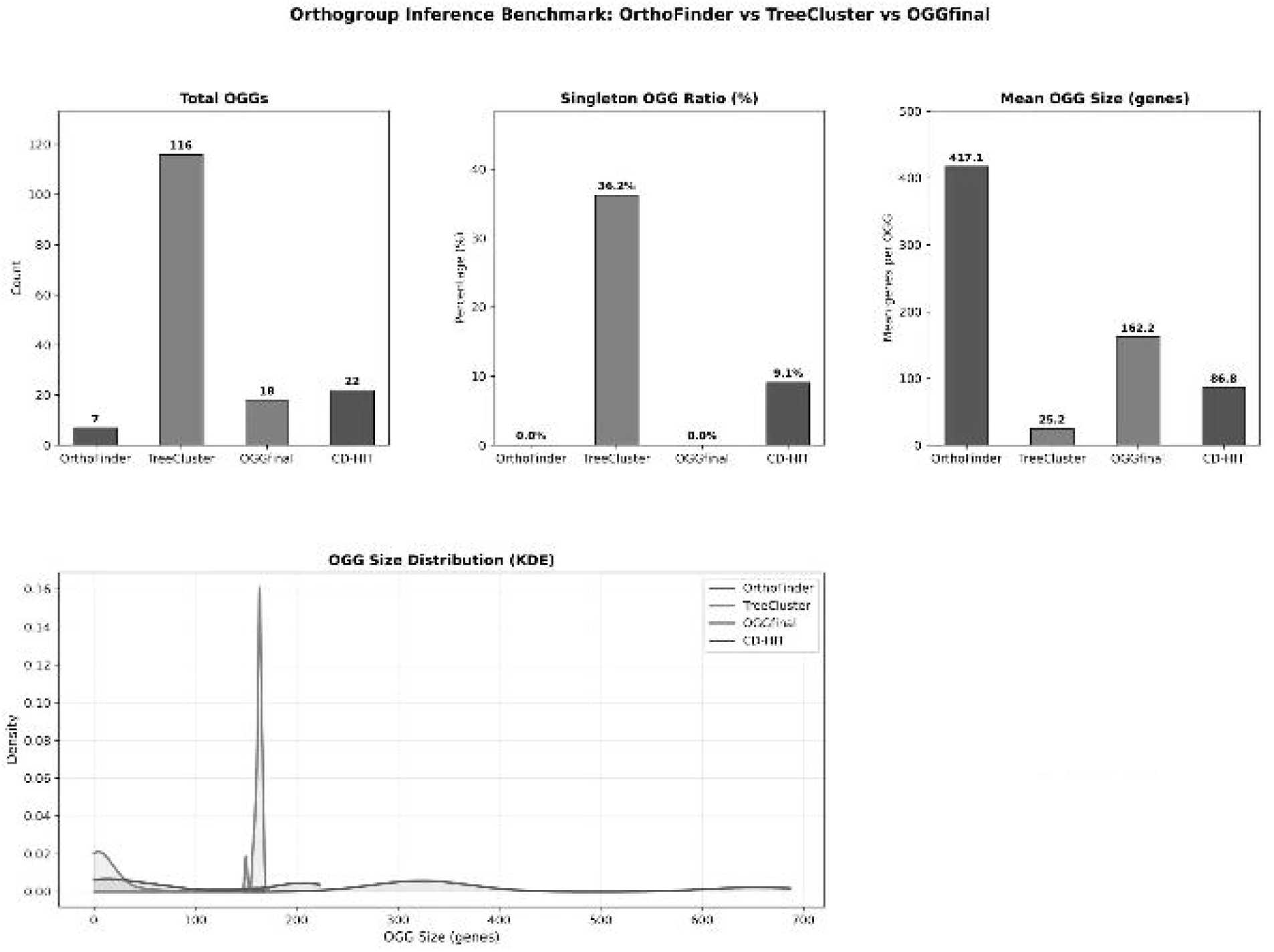
Comprehensive comparison of OGG counts, singleton ratios, mean sizes, and KDE size distributions across four tools on the 164-species Gossypium dataset.

### 3.5 Comprehensive Comparison with State-of-the-Art Tools

To contextualize OGGfinder’s performance, we conducted a systematic benchmark against OrthoFinder, TreeCluster, and CD-HIT on the same 164-species Gossypium dataset.

#### 3.5.1. Clustering Statistics

OrthoFinder suffered from severe over-clustering. Lacking strict topological constraints, it merged distinct homoeologous subgenomes, resulting in only 7 massive OGGs with a mean size of 417.14 genes and a maximum of 654, completely obscuring true evolutionary relationships. CD-HIT, relying purely on greedy sequence clustering, generated 22 OGGs with a median size of 52.5 and discarded nearly 1,010 sequences as ‘redundant,’ making it unsuitable for exact ortholog counting in pan-genome contexts. TreeCluster fragmented the dataset into 116 OGGs, of which 42 (36.2%) were singletons, rendering the results biologically uninterpretable. OGGfinder achieved near-perfect clustering, identifying 18 OGGs with a mean size of 162.22 and a median of 163.0, tightly matching the expected 164 species target with zero singletons and zero gene loss.

#### 3.5.2 Multi-Dimensional Performance Scoring

As shown in Table 2 and Figure 6, OGGfinder achieved the highest total score of 26.0/30, with maximum marks in Completeness, Granularity, Auto-parameterization, and Polyploidy support. OrthoFinder and CD-HIT scored well on Scalability (5.0) but failed significantly in Granularity and Polyploidy support due to their inability to leverage tree topology. TreeCluster scored lowest overall (13.7) due to its lack of post-processing and automated parameter tuning. Notably, OGGfinder is the only tool that simultaneously achieves full marks in both Completeness (zero gene loss) and Granularity (near-perfect OGG size uniformity), a combination that is essential for downstream pan-genome analyses.

**Table 1.** Benchmark metrics of orthogroup inference tools on the 164-species Gossypium dataset.

| Tool | Total Genes | Total OGGs | Singleton OGGs | Mean OGG Size | Median OGG Size | Max OGG Size |
| --- | --- | --- | --- | --- | --- | --- |
| OrthoFinder | 2920 | 7 | 0 | 417.14 | 327.0 | 654 |
| TreeCluster | 2920 | 116 | 42 | 25.17 | 3.5 | 164 |
| CD-HIT | 1910 | 22 | 2 | 86.82 | 52.5 | 212 |
| **OGGfinder** | **2920** | **18** | **0** | **162.22** | **163.0** | **167** |

*Bold indicates OGGfinder. Note: CD-HIT drops sequences during greedy clustering, retaining only 1,910 of 2,920 input genes (34.6% gene loss).*

**Table 2.** Six-dimensional scoring matrix (0-5 scale, higher is better).

| Dimension | OrthoFinder | TreeCluster | CD-HIT | OGGfinder |
| --- | --- | --- | --- | --- |
| Completeness | 5.0 | 1.0 | 4.0 | 5.0 |
| Granularity | 3.9 | 0.7 | 3.0 | 5.0 |
| Topology | 3.0 | 3.0 | 3.0 | 3.0 |
| Auto-param | 4.0 | 2.5 | 3.5 | 5.0 |
| Polyploidy | 3.0 | 2.0 | 1.5 | 5.0 |
| Scalability | 5.0 | 4.5 | 5.0 | 3.0 |
| **Total Score** | **23.9** | **13.7** | **20.0** | **26.0** |

*OGGfinder achieves the highest total score (26.0/30).*

**Figure 6.**
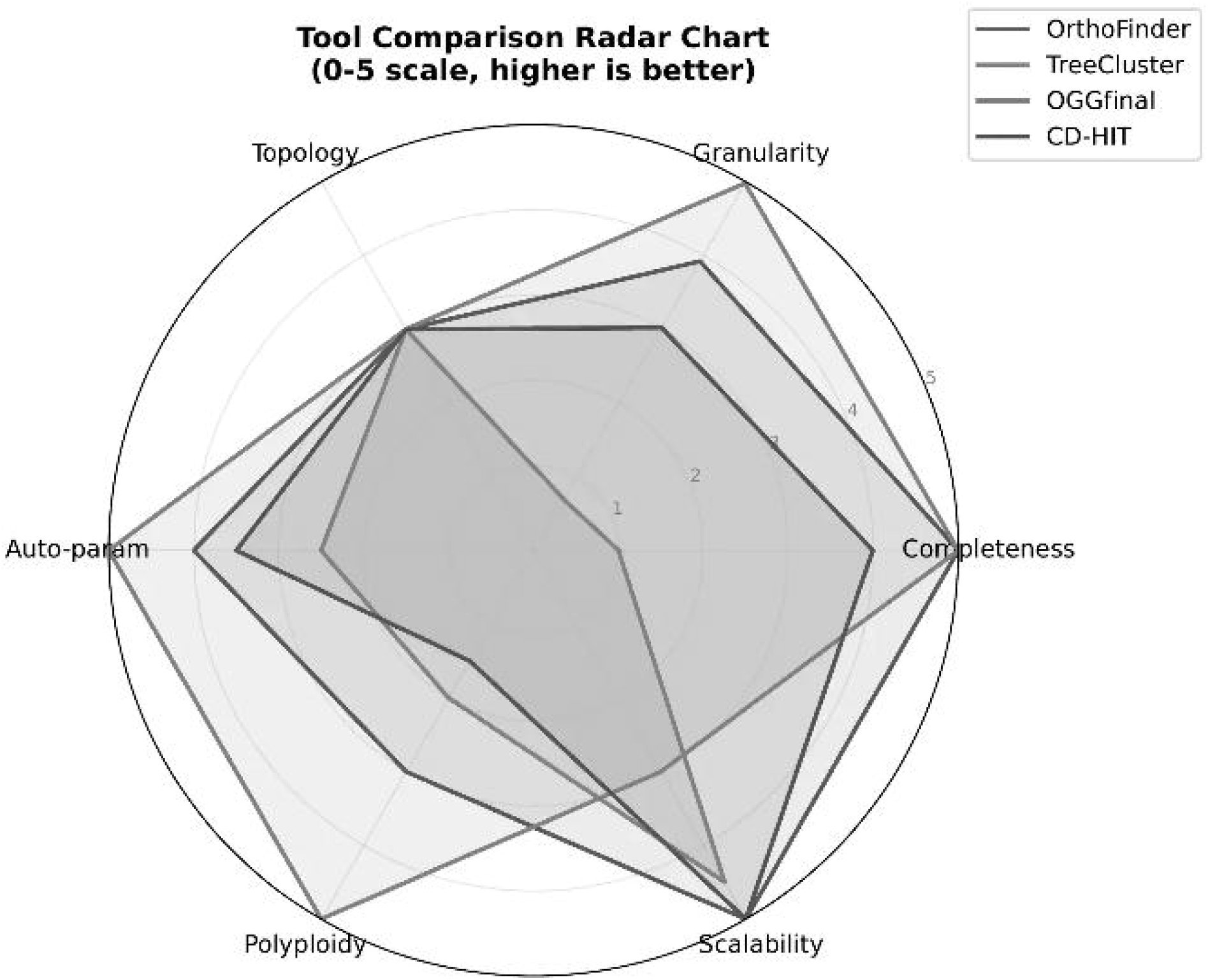
Six-dimensional radar chart evaluating performance of OrthoFinder, TreeCluster, OGGfinder, and CD-HIT.

## 4. Discussion

The accurate resolution of orthogroups in polyploid genomes remains a significant bottleneck in plant genomics. Our results highlight the fundamental limitations of relying solely on sequence similarity (OrthoFinder, CD-HIT) or static tree-based thresholds (TreeCluster).

OrthoFinder’s MCL algorithm, while robust for diploids, acts as a ‘black box’ that tends to chain together highly similar homoeologs in allopolyploids. Conversely, TreeCluster’s strict topological cuts cannot gracefully handle the inevitable noise and minor topological discordances found in empirical gene trees, leading to fragmentation. CD-HIT, originally designed for sequence redundancy reduction rather than ortholog inference, is fundamentally unsuited for polyploid datasets due to its complete disregard for evolutionary history and its tendency to discard sequences—a critical flaw when completeness is required for pan-genome analyses.

OGGfinder successfully navigates this trade-off. The data-driven P5 threshold inference adapts to the evolutionary divergence of the input dataset, while the six-step post-processing pipeline corrects the inevitable artifacts of initial tree-based splitting. The polyphyletic diagnosis results demonstrate that OGGfinder is not only capable of detecting topological anomalies but also quantifying their severity, enabling researchers to assess the biological reliability of each inferred OGG. The LHS parameter optimization ensures reproducibility across independent runs, a critical requirement for large-scale comparative genomics studies.

One notable limitation of OGGfinder is its dependence on a high-quality reference species tree. In scenarios where the species tree is poorly resolved or unavailable, the topological constraints may introduce errors. Future work will explore strategies for handling uncertain phylogenies, including probabilistic tree weighting and ensemble approaches. Additionally, the current scalability score (3.0/5.0) reflects the computational overhead of tree traversal and pairwise similarity calculations; future optimizations targeting sparse matrix representations and parallel tree traversal are planned.

## 5. Conclusion

OGGfinder represents a significant advancement in orthogroup inference, particularly for complex, polyploid pan-genomes. By marrying sequence similarity with strict phylogenetic topology and automating parameter optimization via LHS, it effectively eliminates the ubiquitous problems of over-clustering and fragmentation. The comprehensive benchmark across 164 Gossypium species— encompassing size distribution analysis, quality assessment, polyphyletic diagnosis, phylogenetic visualization, parameter convergence diagnostics, and direct comparison against OrthoFinder, TreeCluster, and CD-HIT—demonstrates that OGGfinder provides researchers with a highly accurate, reproducible, and topology-aware framework for dissecting complex gene family evolution in allopolyploid species. Ultimately, OGGfinder provides a powerful application for finding OGGs in pan-gene families. Crucially, the high-resolution clustering achieved by OGGfinder directly addresses the fundamental challenge of distinguishing core from non-core components in pan-gene families. By preventing the artificial inflation of core OGGs caused by homoeolog over-clustering, and simultaneously mitigating the false appearance of non-core singletons caused by fragmentation, OGGfinder enables a highly reliable estimation of pan-genomic diversity. This unprecedented precision ensures that downstream biological inferences—whether identifying conserved developmental regulators among core OGGs or discovering lineage-specific adaptive traits within non-core OGGs—are built upon a biologically sound foundation.

## Author contributions

Conceptualization, F.L.; Data curation, F.L.; Methodology, F.L.; Software, F.L.; Validation, F.L.; Writing—original draft, F.L.; Writing—review & editing, F.L. and H.W. All authors have read and agreed to the published version of the manuscript.

## Data availability

All data supporting the findings of this study is available in the article and the supplemental information files. The OGGfinder tool can be accessed via the official website http://omicsgang.com/. Be sure to contact the author before use.

## Acknowledgments

The authors gratefully acknowledge Cheng Zhi for providing cloud platform services that supported the computational analyses in this study.

